# The Human Telomeric Proteome During Telomere Replication

**DOI:** 10.1101/2020.06.14.150524

**Authors:** Chih-Yi Gabriela Lin, Anna Christina Näger, Thomas Lunardi, Aleksandra Vančevska, Gérald Lossaint, Joachim Lingner

**Author notes:** Correspondence (J.L.) and (G.L.).

## Abstract

Telomere shortening can cause detrimental diseases and contribute to aging. It occurs due to the end replication problem in cells lacking telomerase. Furthermore, recent studies revealed that telomere shortening can be attributed to difficulties of the semi-conservative DNA replication machinery to replicate through the bulk of telomeric DNA repeats. To investigate telomere replication in a comprehensive manner, we develop QTIP-iPOND, which enables purification of proteins that associate with telomeres specifically during replication. We identify in addition to the core replisome a large number of proteins that associate with telomere replication forks and validate their importance. We find that POT1 is depleted, whereas histone H1 is specifically enriched, at telomere replication forks. Our work reveals the dynamic changes of the telomeric proteome during replication, providing a valuable resource of telomere replication proteins. To our knowledge, this is the first study that examines the replisome at a specific region of the genome.

## Introduction

Telomeres are crucial for maintaining chromosome structure and genome stability. In humans, telomeric DNA consists of 5’-TTAGGG-3’/5’-CCCTAA-3’ repeats with an overall length of 5,000 to 15,000 nucleotides. Telomeric DNA recruits a large number of proteins, which provide telomere functions at chromosome ends. Most prominent is the six subunits containing shelterin protein complex: TRF1 (telomere repeat binding factor 1), TRF2 (telomere repeat binding factor 2), POT1 (protection of telomeres 1), TPP1 and TIN2 (TRF1-interacting nuclear factor 2)^1^. In addition to the shelterin components, a large number of additional proteins associates with telomeres^2–4^. Experimental manipulation and genetic data linking mutant gene products to telomere syndromes have revealed important functions for several of these proteins.

Telomere shortening in telomerase-negative somatic cells triggers a DNA damage response^5^. This leads to cellular senescence providing a powerful tumor-suppressive mechanism as it limits the proliferation of cells in precancerous lesions^6^. On the other hand, premature telomere shortening can lead to a number of degenerative disorders giving rise to bone marrow failure, pulmonary fibrosis and cancer^7–11^. In these telomere syndromes, telomere maintenance by either semi-conservative DNA replication or telomerase is defective.

Telomeres pose the following challenges to the DNA replication machinery. First, unwinding of the parental DNA during telomere replication exposes the G-rich single stranded DNA which can form highly stable G-quadruplex (G4) structures. G4 structures can cause stalling or collapse of replication forks, requiring specialized helicases including WRN, RTEL1 and BLM for unwinding^12–15^. Second, the t-loop in which the telomeric 3’ overhang is tucked into the double stranded part of the telomeric DNA^16^ needs to be unwound during telomere replication by the RTEL1 helicase^17–19^. Third, the long noncoding RNA TERRA is transcribed from a large number of chromosome ends^20–23^. TERRA can form RNA/DNA hybrid structures which interfere with DNA replication if not removed by RNase H, the THO complex, RNA surveillance factors or the splicing factors SFPQ and NONO^24–28^. Finally, replication origins are present in the subtelomeric regions but replication initiates only rarely within the telomeric repeats^14,29^. Therefore, replication at telomeres is mostly unidirectional and stalled forks may not be rescued from converging forks that come from the end of chromosomes. In order to specifically characterize the replication machinery that overcomes the many hurdles posed at telomeres, we combine QTIP (quantitative telomeric chromatin isolation protocol)^4,30^ to purify telomeric chromatin with iPOND (isolation of proteins on nascent DNA)^31,32^ which enriches proteins associated with replicating DNA. We identify a large set of proteins that are specifically enriched at telomeres during their replication. Crucial functions for several of these proteins are demonstrated. Moreover, we find that the single strand telomeric DNA binding POT1 is depleted from replication forks suggesting that it must be replaced by RPA for telomere replication. On the other hand, histone H1 variants are present during telomere replication and are depleted from telomeres only later when telomeric chromatin matures. This study provides comprehensive insights into the dynamic changes of telomeric proteins during telomere replication.

## Results

### QTIP-iPOND enables purification of telomeric chromatin during replication

To examine the dynamic changes of the telomeric proteome that occur during replication, we developed QTIP-iPOND, which combines two previously established techniques in a consecutive manner: In the first step, crosslinked telomeric chromatin is purified by QTIP using antibodies against TRF1 and TRF2^4,30^. In the second step, the purified telomeric chromatin is further fractionated by iPOND^31,32^ to separate the telomeric chromatin present at DNA replication forks from non-replicating telomeric chromatin (Figure 1a). We took advantage of HEK293E suspension cells to expand large scale culture. Moreover, we endogenously FLAG-tagged TRF1 and TRF2 in HEK293E cells. This allowed immunoprecipitation of telomeric chromatin during QTIP purification with anti-FLAG antibodies and elution of purified telomeric chromatin with excess FLAG peptide without denaturing protein structures. To label nascent DNAs at replication forks, we incubated cells with EdU (5-Ethynyl-2’-deoxyuridine), a thymidine (Thy) analog for 10 minutes. In the EdU pulse experiment, cells were fixed directly after EdU incubation. EdU-labeled nascent DNA was further covalently linked with biotin-azide using a click reaction. After cell lysis, we first purified telomeric chromatin using anti-FLAG beads targeting the FLAG-tagged TRF1 and TRF2. Telomeric chromatin was eluted with excess FLAG-tag peptide, which competed off the telomeric chromatin from anti-FLAG beads. In the second step, the telomeric chromatin associated with nascent replicating DNA was separated from non-replicating telomeric chromatin using streptavidin beads targeting the biotinylated EdU-labeled DNA. Moreover, we carried out a Thy chase experiment in which the EdU-containing medium was replaced by Thy-containing medium after EdU incorporation and was harvested 4 hours after Thy addition. (Figure S1a; see STAR Methods for details).

**Figure 1.**
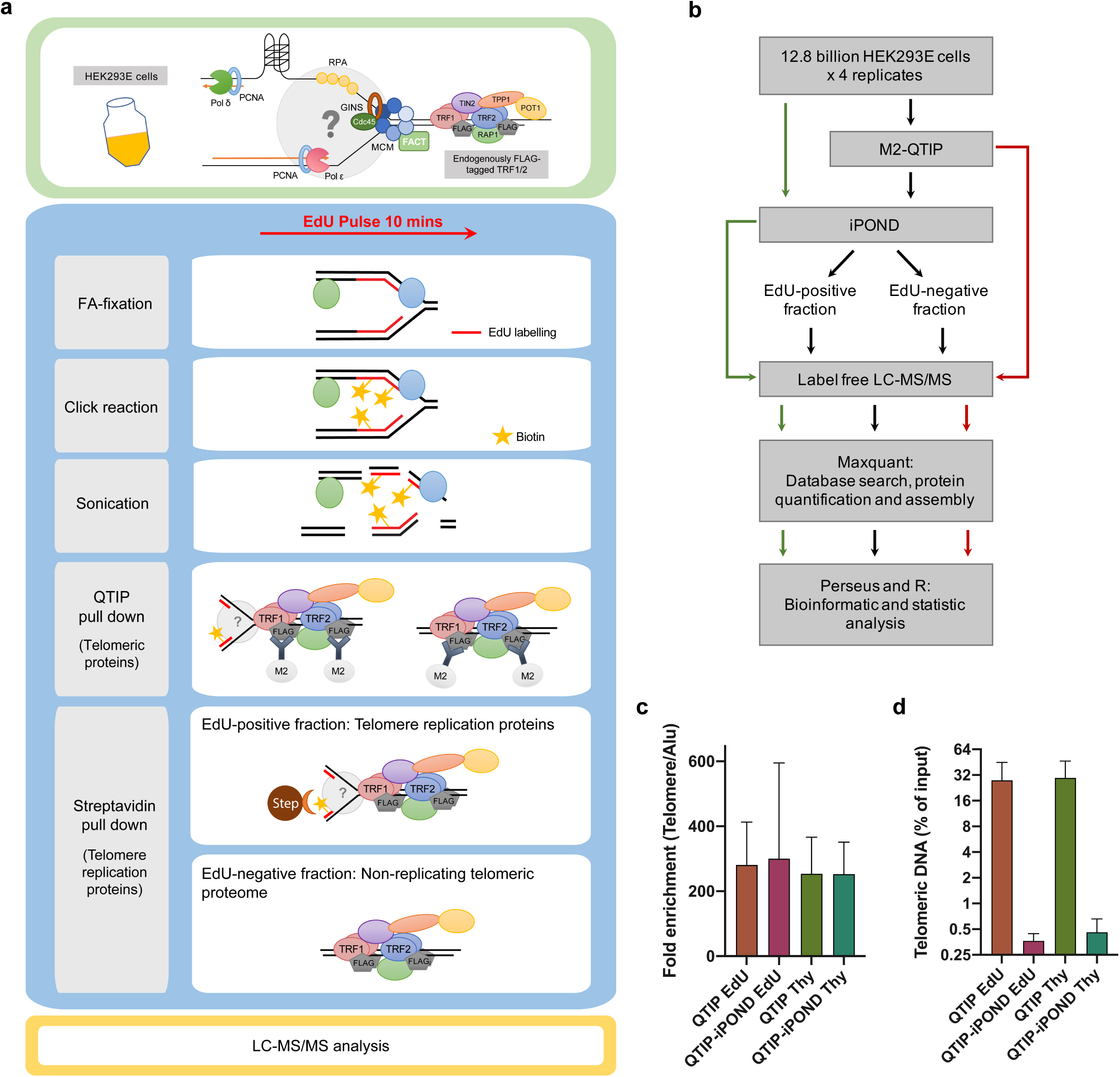
Experimental scheme of QTIP-iPOND EdU pulse. a. To identify proteins near telomere replication forks, we combined two techniques: the QTIP (quantitative telomeric chromatin isolation protocol) to identify proteins present at telomeres and the iPOND (identification of proteins on nascent DNA) to identify proteins present at DNA replication forks. FA: formaldehyde; Strep: streptavidin. b. Flow chart of QTIP-iPOND experiment. 12.8 billion cells were used for each replicate, which was analyzed by QTIP-iPOND (9.6 billion cells for EdU pulse and Thy chase), iPOND (1.6 billion cells for EdU pulse and Thy chase) and QTIP (1.6 billion cells for FLAG-tagged TRF1/2 HEK293E and non-tagged WT HEK293E. c. Telomeric DNA recovery (5 biological replicates; 3 of them are from the full-scale experiment). d. Fold enrichment of telomeric DNA over Alu element (4 biological replicates; 2 of them are from the full-scale experiment). Data are represented as mean ± SD. (See also Figure S1a and S1b)

Four biological replicates, in which a total of 12.8 billion cells were used each time, were performed. Each replicate was analyzed by QTIP-iPOND, iPOND and QTIP. For the QTIP-iPOND analyses, the EdU-positive fraction [EdU(+)], which contained telomeric chromatin during replication, and the EdU-negative fraction [EdU(-)], which contained non-replicating telomeric chromatin, were analyzed using LC-MS/MS (Figure 1b). To evaluate the enrichment and recovery of telomeric chromatin in the QTIP and QTIP-iPOND samples, we performed dot blot hybridization with a telomeric probe and a probe of Alu elements, representing non-telomeric regions. Telomeric DNA was enriched approximately 250-fold over Alu element-containing DNA (Figure 1c) and the recovery of telomeric DNA with QTIP was around 30%. The average yield of telomeric DNA in the QTIP-iPOND samples (0.3-0.4%) was around 100-fold lower than in the QTIP samples (Figure 1d). The low final yield of telomeric DNA in QTIP-iPOND chromatin was expected because the EdU pulse lasted only 10 minutes, only 18% of the 293E cells were in S phase upon harvest (Figure S1b), and replication of one telomere may last only a couple of minutes. However, the large-scale experiments gave sufficient material to allow analysis by mass spectrometry.

### Proteomic analysis reveals proteins that are enriched at telomeres during DNA replication

In order to identify the proteins enriched near telomeric replication forks, we compared the intensity of proteins present in EdU-positive versus EdU-negative fractions [EdU(+)/EdU(-)] obtained from the EdU pulse labeled cells using label-free quantification (LFQ) (Figure 1). Since telomeric proteins were purified by pulling down TRF1 and TRF2, and TRF1 is present at non-replicating as well replicating telomeres^29^, we normalized protein amounts to TRF1. 1,732 nuclear proteins were identified. We first examined 40 core replication proteins which have known functions for genome-wide DNA replication and were identified here as well as in several replisome purification approaches before ^33^. 35 of the core replication proteins were enriched more than 2.8 fold in EdU(+)/EdU(-) telomeric chromatin fractions (Figure 2a). These enrichment factors are comparable to the ones reported for iPOND samples^32,34,35^, indicating the successful enrichment of replication proteins in the EdU(+) fractions upon QTIP-iPOND purification.

**Figure 2.**
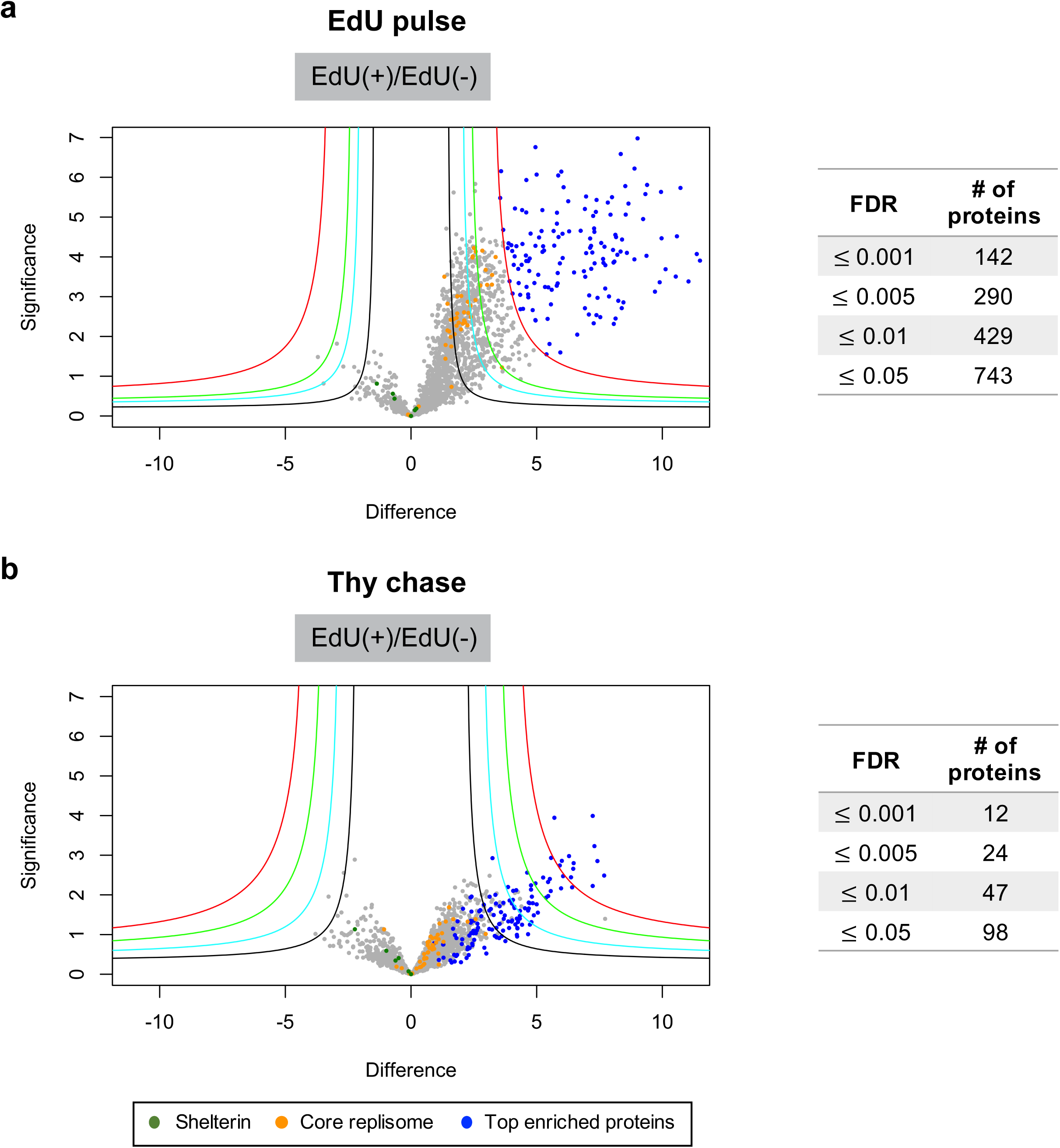
Protein distribution of QTIP-iPOND. a. EdU positive over EdU negative samples in the EdU pulse experiment. b. EdU positive over EdU negative samples in the Thy chase experiment. Difference: log_2_(Fold change); Significance: −log(p-value); Blue circles: the 142 proteins which FDR ≤ 0.001; Orange circles: core replication proteins; Dark green circles: shelterin components; Red curves: FDR=0.001; Green curves: FDR=0.005; Cyan curves: FDR=0.01; s0 = 3. (See also Figure S2 and Table S1)

Next, we determined which proteins were more than 2.8-fold (log_2_(1.5)) enriched in EdU(+)/EdU(-) and had an FDR (false discovery rate) ≤ 0.05 (Figure 2a and Table S1). 743 proteins fulfilled these criteria of which 142 had an FDR ≤ 0.001. The enrichment of these proteins was also strongly reduced in the Thy chase EdU(+) samples (Figure 2B), confirming that these proteins were enriched at telomeres during their replication. Of note, in the Thy chase experiments, we observed a slow recovery of telomere replication which may have been due to the cellular stress that was induced by repeated pelleting and resuspension of cells before thymidine incubation. In support of a slower recovery of replication at telomeres versus the bulk of the genome after Thy addition, we observed a 3.2-fold enrichment of PCNA in EdU pulse/Thy chase in iPOND samples against only a 1.9-fold enrichment in the QTIP-iPOND chromatin (Tables S1 and S2). Therefore, the majority of replication proteins and the 142 top enriched proteins were still enriched at telomeres in the EdU(+) QTIP-iPOND fraction even after the chase, but to a much lesser extent (Figure S2a). As expected, the EdU-negative fractions obtained in QTIP-iPOND, which mainly contained the non-replicating telomeric proteome, showed no striking differences between the EdU pulse and the Thy chase (Figure S1b and 3b).

### Core replisome components detection at telomeres by QTIP-iPOND

In addition to QTIP-iPOND, we simultaneously performed iPOND analyses during EdU pulse and Thy chase in order to distinguish proteins that are specifically present at telomere replication forks from general chromatin bound (Figure 1b). 344 proteins were enriched in iPOND during the EdU pulse. Of these, 105 proteins were enriched in both iPOND and QTIP-iPOND (Figure S3a and Table S2). 40 proteins that represent core replication proteins are listed in Figure 3, including RPA subunits, PCNA, MCM proteins, RFC proteins, FACT complex, MSH proteins, DNA polymerases and others. The enrichment of these proteins was similar in all four biological replicates of QTIP-iPOND, indicating that the enrichment of these proteins at telomere replication forks was reproducible and significant.

**Figure 3.**
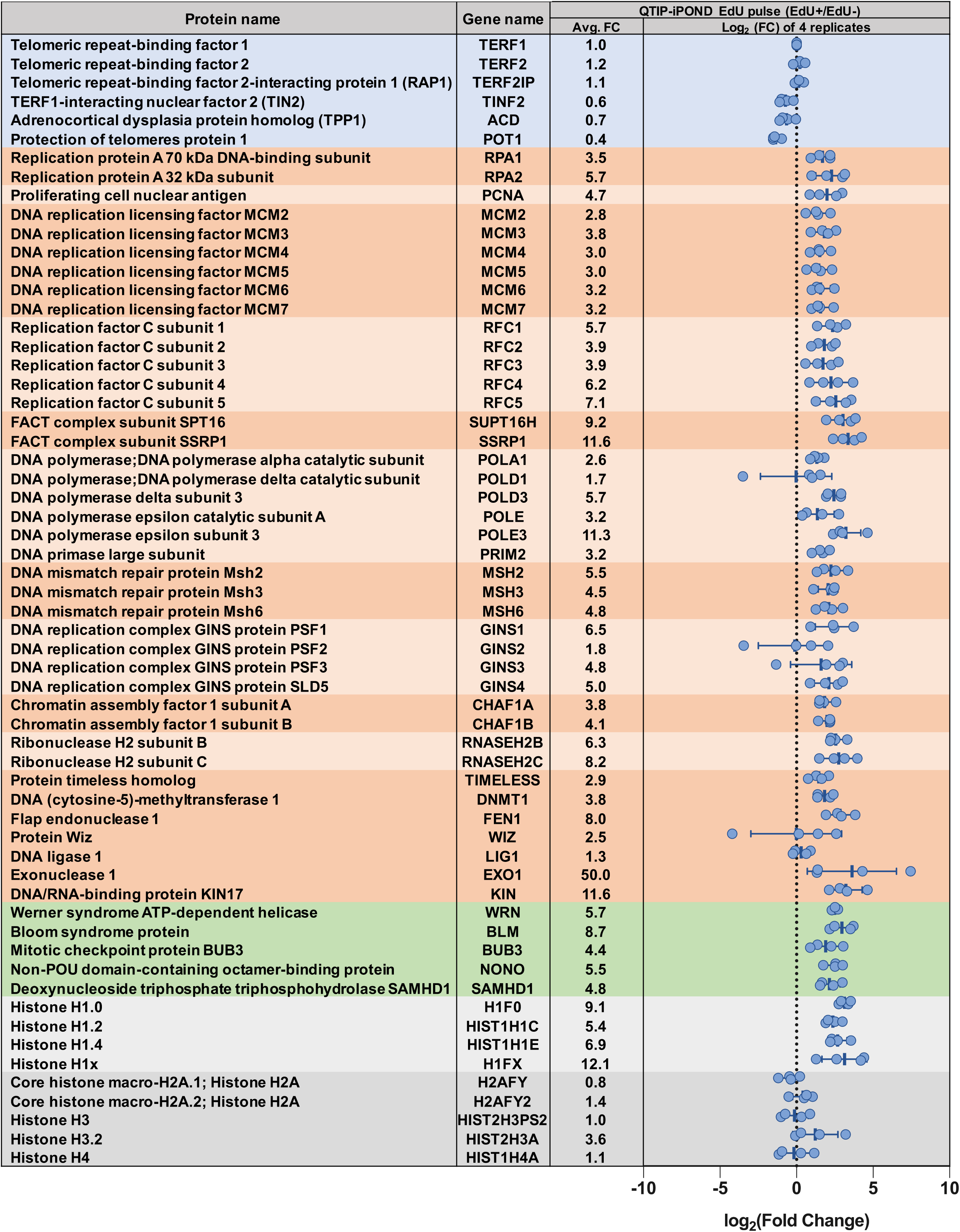
List of proteins present in the QTIP-iPOND experiment. Proteins in the blue background are shelterin components; proteins in the dark/light orange background are core replication proteins; proteins in the green background are telomere replication proteins; proteins in the dark/light grey are histones. Each blue circle indicates the log_2_(Fold Change) in one replicate. Avg. FC: the average fold change. Data are represented as mean ± SD.

Moreover, we identified proteins that had been shown to function specifically in telomere replication (Figure 3). This includes WRN, BLM, NONO and SFPQ. We also identified BUB3, which has been shown to promote telomeric DNA replication in human cell lines ^36^. SAMHD1 was also enriched at telomere replication forks. It has been recently identified by QTIP in oncogene-transformed human lung fibroblasts (HLFs) and is important for telomere integrity upon TRF1 depletion^37^. RTEL1 was not detected in both QTIP-iPOND and QTIP (Table S1 and S3), possibly due to its low abundance at telomeres. Overall, these results indicated that telomeric replication proteins were specifically purified by QTIP-iPOND.

### Histone H1 variants are present during telomere replication

In previous iPOND experiments, it has been shown that the levels of core histone proteins remain unchanged during replication^32,38^. Consistently, our QTIP-iPOND analyses indicated that histones H2A, H3 and H4 are similarly abundant in replicating EdU(+) and non-replicating EdU(-) telomeres (Figure 3). However, one of the replicative variants, histone H3.2 was enriched in the EdU(+) fraction in two of the replicates. Furthermore, to our surprise, we identified that four histone H1 variants were consistently enriched in the EdU(+) samples (Figure 3). In contrast, H1 variants were depleted in the EdU(+) iPOND samples (Figure S3b), which is consistent with previous iPOND experiments^35,38^.

Previous studies had shown that nucleosomes are unusually narrowly spaced at telomeres^39,40^ and that histone H1 is depleted^3^. In apparent contradiction to these findings, it was observed that the depletion of histone H1 can increase the level of telomere sister chromatid exchange (T-SCE) and telomere length in murine embryonic stem cells^41^. Our result appears to be consistent with these previous observations indicating that histone H1 variants are enriched at telomeres only during replication. Thus, H1 may be important for maintaining telomere stability upon synthesis.

### Telomerase subunits are detected near telomere replication forks

From the 142 top enriched proteins, 139 were specifically enriched in QTIP-iPOND but not in iPOND (Figure 4). Among these proteins were Cajal body (CB) and nucleolar proteins. Interestingly, three of the four H/ACA ribonucleoprotein complex subunits, DKC1, NHP2 and GAR1, which associate with the telomerase RNA moiety hTR, were identified. Although we did not detect hTERT in our MS samples, which may be due to technical limitations, the identification of three telomerase subunits suggests that the telomerase complex or subcomplexes may travel with telomere replication forks as suggested previously but never demonstrated^42,43^. Of note, EdU is not a substrate for telomerase^44^, indicating that in QTIP-iPOND we did not purify telomeric chromatin during elongation by telomerase. Moreover, other nucleolar proteins which have known functions related to telomere biology were identified. This includes PARN, a poly(A)-specific ribonuclease which is involved hTR maturation^45–47^; NOLC1 which interacts with TRF2 and has been reported to regulate nuclear and nucleolar TRF2 localization^48^; nucleostemin which regulates TRF1 dimerization and telomere association^49,50^; GNL3L which stabilizes TRF1^51^; PINX1, a Pin2/TRF1-interacting protein which mediates the accumulation of TRF1 in nucleoli and also interacts with TRF1 in the nucleoplasm enhancing TRF1 binding to telomeres during S phase^52^. The recruitment of these CB and nucleolar proteins suggests interactions between these compartments during telomere replication.

**Figure 4.**
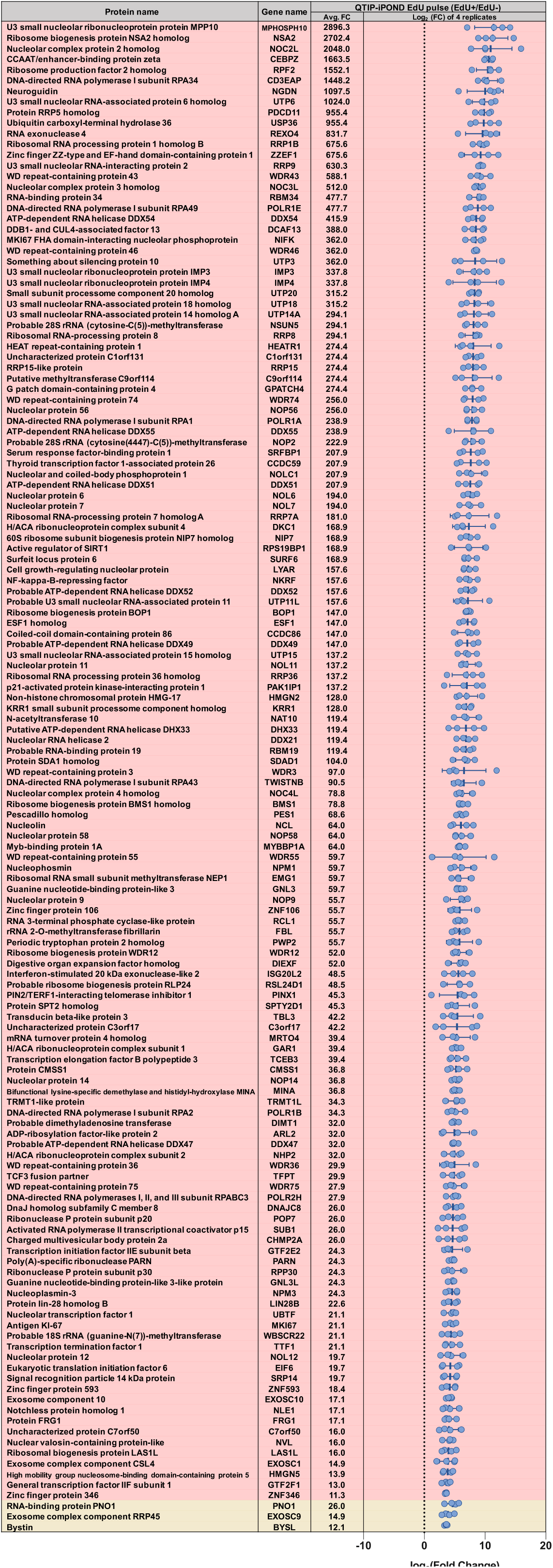
List of the 142 top enriched proteins. The list indicates the 142 proteins for which the FDR is below 0.01. Each blue circle indicates the log_2_(Fold Change) in one replicate. Proteins in the red background are specifically enriched in telomere replication; proteins in the light yellow background are enriched in both QTIP-iPOND EdU pulse and iPOND EdU pulse versus Thy chase. Avg. FC: the average fold change. Data are represented as mean ± SD.

### Telomere fragility assay reveals the requirement of novel proteins identified by QTIP-iPOND for efficient telomere replication

To assess the functional relevance of QTIP-iPOND enriched proteins for telomere replication, we chose 24 QTIP-iPOND enriched proteins that were not present in iPOND only fractions and had not been previously linked to telomere biology. We depleted these proteins from HeLa-Long cells using siRNA pools (Figure 5 and Figure S4). After siRNA transfection, telomere morphology of metaphase chromosomes was analyzed by fluorescence *in situ* hybridization with a telomere specific probe (Figure 5a). Telomeres that showed smeary or multiple telomeric signals upon gene depletion were recognized as fragile telomeres and inferred to have telomere replication defects^29^. Apart from the positive control siTRF1, we depleted five additional known telomere replication proteins (BLM, WRN, NONO and SAMHD1) that were present in our QTIP-iPOND samples. Out of the 29 depleted proteins, 17 gave a telomere fragility score that was higher than 0.4 (high fragility), 5 of them scored between 0.2 to 0.4 (moderate fragility), and 7 of them were scored below 0.2 (low fragility) in the first experiment (Figure 5b and Figure S4). We repeated the fragility analysis for the 17 proteins that had higher scores than 0.4. 13 of them remained in the same high fragility category (score ≥ 0.4) in experiments carried out in triplicate, which includes 9 novel telomere replication proteins and all 4 positive controls (Figure 5b-5d). The depletion levels of the mRNAs for all 17 proteins were determined by RT-qPCR, showing a reduction of mRNA levels ranging from 54 to 95% (Figure S5). Altogether, our results support the notion that a large fraction of proteins identified by QTIP-iPOND protects telomeres from fragility, indicating that they sustain telomere replication.

**Figure 5.**
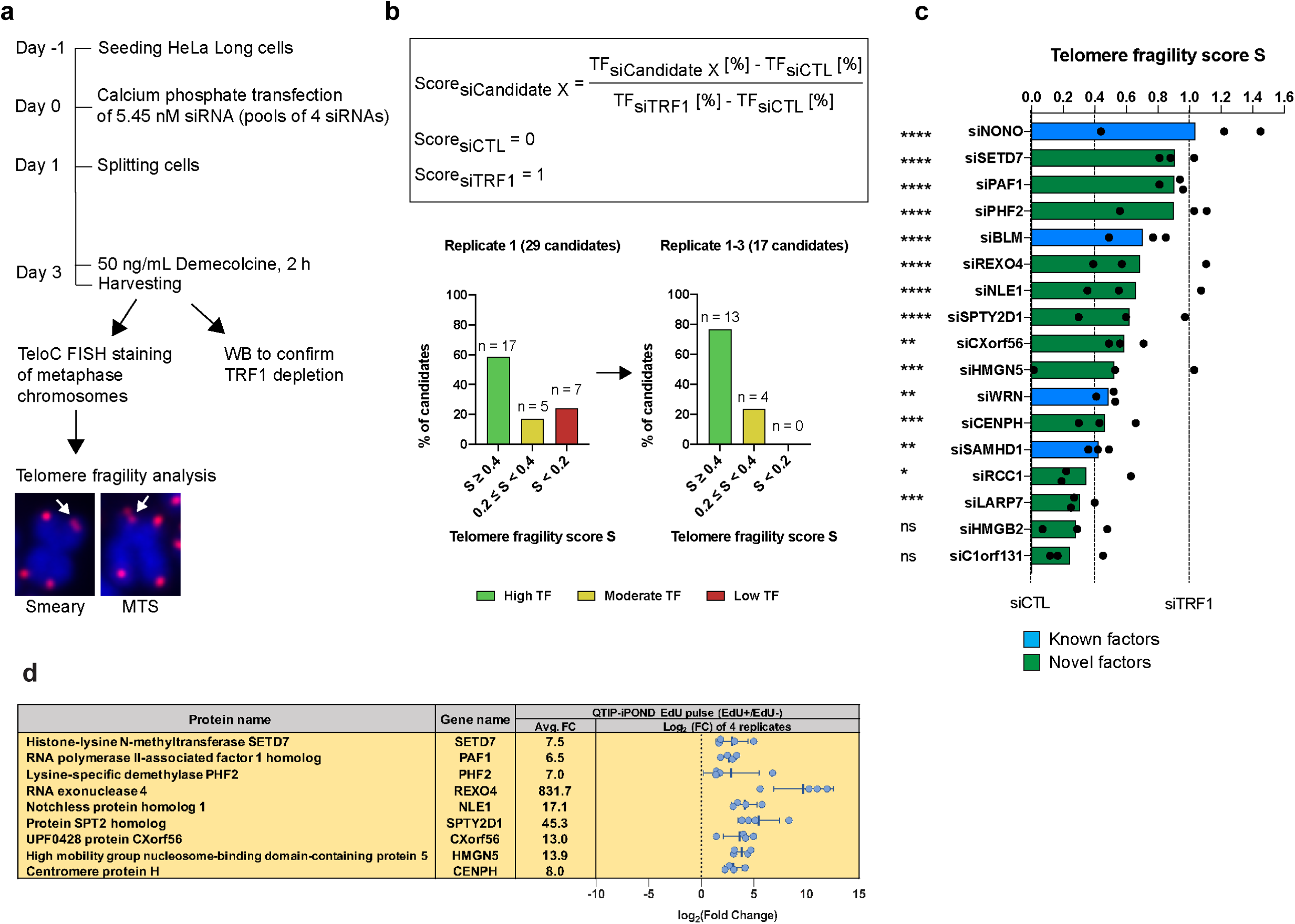
Candidate validation through telomere fragility analysis. a. Experimental outline of siRNA screen for telomere fragility using HeLa-Long cells (with an average telomere length of 33 kb). MTS: multiple telomeric signals. Pixel size of FISH images was reduced 4-fold for illustration. b. Percentage of tested candidates with respective telomere fragility score. The percentage of fragile telomeres per metaphase was determined for each depleted protein candidate and compared to TRF1-depleted cells (siTRF1) and control transfected cells that had been transfected with a non-targeting siRNA (siCTL). A score of 1 corresponds to the amount of telomere fragility detected in TRF1-depleted cells and a score of 0 to the amount of telomere fragility detected in the negative control (siCTL). TF: telomere fragility c. Telomere fragility score for 3 biological replicates with 45 metaphases analyzed per condition in total. Statistical analysis was done by unpaired t-test of telomere fragility per metaphase compared to siCTL. * < 0.05, ** < 0.01, *** < 0.001, **** < 0.0001. Blue and green indicate known and novel factors that prevent telomere fragility, respectively. d. QTIP-iPOND EdU pulse for novel factors with average telomere fragility score > 0.4. Data are represented as mean ± SD. (See also Figure S4 and S5)

### POT1 is depleted from replication forks

The abundance of proteins in replicating EdU(+) versus non-replicating EdU(-) telomeric chromatin had been normalized to TRF1. Thus, it was not unexpected that the shelterin components TRF2 and Rap1 did not change in abundance relative to TRF1 (Figure 3). However, TIN2, TPP1 and most noticeably POT1 were less abundant in the replicating versus non-replicating telomeric chromatin fractions, with POT1 being reduced by 60% (Figure 3). To confirm that POT1 was reduced at replication forks, we pulse labeled HEK293E cells with BrdU for 10 or 30 minutes which respectively labeled telomeres near replication forks or preferentially telomeres that had been fully replicated (Figure 6a). We avoided to perform the chase experiment to ensure that the replication was uninterrupted. Crosslinked chromatin was immunoprecipitated (ChIP) using antibodies against POT1, TRF1 and RPA32. The immunoprecipitated DNA was analyzed with an anti-BrdU antibody to infer association of these proteins with BrdU-labeled DNA and hybridized with a telomeric probe to reveal telomere association (Figure 6). We compared BrdU intensities between 30-min BrdU incorporation and 10-min BrdU incorporation normalized to the amounts of telomeric DNA. The obtained relative amounts of BrdU were normalized to the ones obtained with the TRF1-ChIP. We observed that the relative BrdU amounts obtained with the POT1-ChIP increased 2-fold with increased time of BrdU incorporation when compared to TRF1. This indicates that POT1 is underrepresented at replication forks (Figure 6c). Notably, the BrdU amounts obtained with the POT1-ChIP were at a similar level as the negative control IgG-ChIP after 10-min of BrdU incubation, indicating very low POT1 abundance at replication forks (Figure S6). In contrast to POT1, the RPA32-ChIP showed a 2-fold decrease of the amounts of BrdU at the 30-min BrdU time point in comparison to the TRF1-ChIP, indicating that RPA32 was highly enriched at telomere replication forks. Overall, our results showed that RPA32 is enriched whereas POT1 is strongly depleted near replication forks. Therefore, POT1 associates with telomeres only upon completion of replication upon telomere maturation.

**Figure 6.**
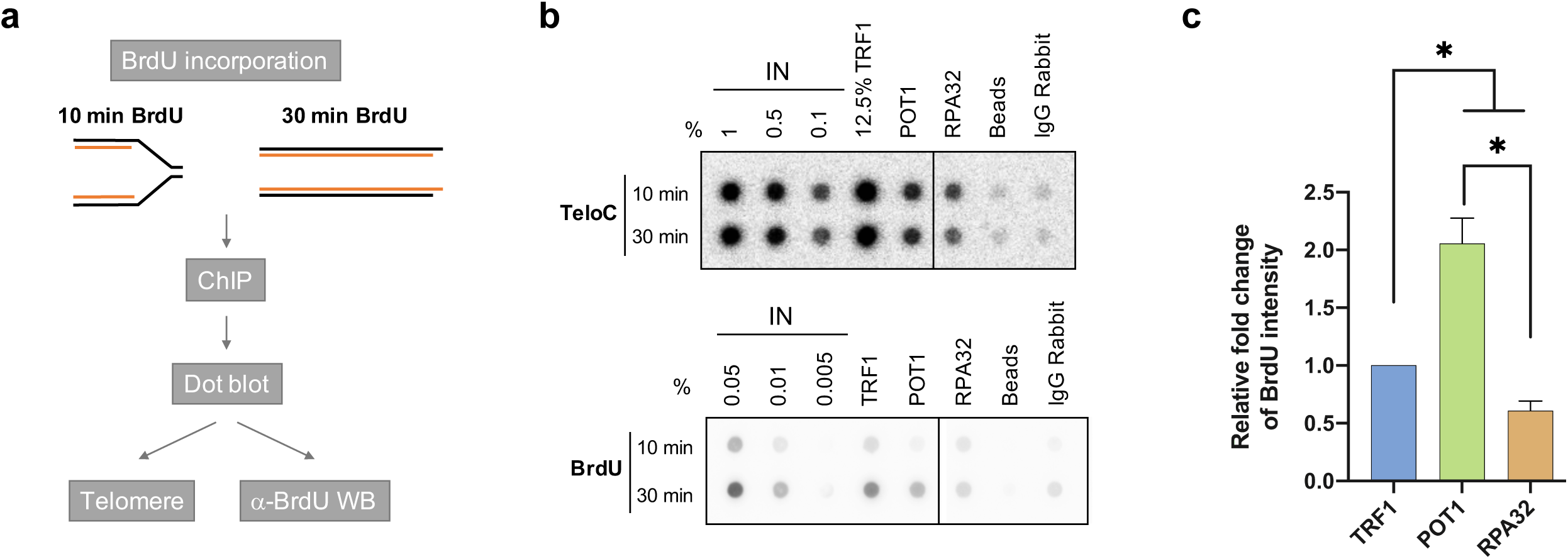
POT1 is depleted from replication forks. a. Flow chart of BrdU-incorporated ChIP. b. DNA pulled-down by ChIP with 10 min or 30 min BrdU incorporation was hybridized with C-strand telomeric probes or was recognized by an anti-BrdU antibody. Beads: Beads only control. c. The relative fold changes of 30-min BrdU intensity and 10-min BrdU intensity from ChIP-dot blot. The results were normalized to the telomere recovery rate and then to TRF1-ChIP. Statistical analysis was done by unpaired t-test to TRF1. * <0.05. Data are in duplicate and are represented as mean ± SD. (See also Figure S6)

## Discussion

The successful replication of telomeres is crucial for maintaining genome stability and preventing cancer and telomere syndromes, yet the molecular mechanisms are not fully understood. Here, we developed QTIP-iPOND to investigate telomere replication in a comprehensive manner. The method allowed the comparison of replicating and non-replicating telomeric proteomes as the telomeric chromatin obtained by the QTIP step was further fractionated into EdU-positive and EdU-negative telomeric fractions using the iPOND protocol. The pulse-chase experiment that was classically employed in iPOND experiments causes pronounced replication stalling at telomeres. Therefore, our design circumnavigated this problem and was able to reveal the dynamic changes of proteins that occur at telomeres during replication.

In our quadruplicate experiment, we identified the majority of core replication proteins and known telomere replication factors using a stringent selection threshold (2.8-fold change and FDR ≤ 0.05). We identified 743 proteins near telomere replication forks of which 638 were specifically present at telomeres but not at non-telomeric replication forks. Of the 638 proteins, we validated 29 proteins using telomere fragility analysis. 76% of them showed moderate to high telomere fragility when depleted. Nine proteins that showed strong telomere fragility upon depletion had not been linked to replication before. Moreover, we identified 142 proteins which were very highly and significantly enriched near telomere replication forks (FDR <0.001), of which only three were detected by iPOND. This supports the notion that the telomeric replisome requires a large set of specialized factors. We also identified telomerase subunits enriched in the replicating telomeric proteome, suggesting that the telomerase complex is present. Life cell imaging has demonstrated that telomerase makes during S phase thousands of transient telomere interactions that last less than one second^53^. Longer lasting interactions occur only when telomerase engages with the telomeric 3’ overhang during telomere elongation. However, we expect that active telomerase-mediated telomere elongation is not captured by QTIP-iPOND as EdU inhibits telomerase *in vitro*^44^. Thus, it seems that the detection of telomerase subunits in replicating chromatin is due to transient short-lived interactions with telomeres.

Interestingly, among the top enriched proteins were several that preferentially localize to Cajal bodies (CBs) or nucleoli. CBs contain telomerase RNA and associate with telomeres in human cells in late S phase. Moreover, hTERT has been found to associate with CB as well as with nucleoli during S phase. ^54,55^. Many nucleolar proteins such as PINX1, PARN, nucleostemin and GNL3L have also been reported to interact with shelterin components and participate in the regulation of telomere length^45,46,51,52,56^. Furthermore, six of the nine novel telomere replication proteins that were verified in our study (SETD7, PHF2, REXO4, NLE1, SPTY2D1 and CENPH) show nucleolar localization according to the gene ontology resource^57,58^. The detailed mechanisms by which these novel telomere replication factors contribute to telomere maintenance need to be unraveled in the future. Overall, our results suggest cross talks between telomeres, CBs and nucleoli. It is tempting to speculate that these structures contribute to telomere replication by their propensity to undergo phase separation.

Linker histone H1 is the least well understood histone protein^59^. Several studies in different species have suggested that histone H1 participates in regulating the timing of replication, especially at late replicating regions^60,61^. Surprisingly, we observed that histone H1 is enriched at telomeres during their replication, whereas it is depleted from mature telomeric chromatin^3^. Since previous studies showed an increase of T-SCE and an elongation of telomere length in H1-depleted murine embryonic stem cells^41^, our results suggest that histone H1 specifically contributes to telomere stability during replication.

Our data show that POT1 is depleted from replication forks at telomeres. Previous studies suggested roles of POT1 in semi-conservative DNA replication. Overexpression of POT1 mutants which are prevalent in T cell lymphoma in human fibrosarcoma HT1080 cells increased telomere fragility indicative of DNA replication defects^62^. On the other hand, it has been proposed that POT1 is replaced by RPA at telomeres in S phase^63^. Our data support the notion that POT1 is replaced by RPA for telomere replication. Therefore, we suspect that the telomere fragility elicited upon overexpression of POT1 mutants was due to an indirect role of POT1 for telomere maintenance, which may relate to its function in suppressing homology directed repair at telomeres, which in turn promotes telomere fragility^64^.

To conclude, QTIP-iPOND enabled the discovery of the dynamic changes that occur in the telomeric proteome to enable telomere maintenance. QTIP-iPOND not only identified the core components of the replicative machinery, but also the large set of protein components which ensures that telomere replication occurs smoothly at all chromosome ends. Our data provides a comprehensive resource and describes a sophisticated machinery that orchestrates hundreds of proteins to accomplish telomere replication.

## Acknowledgments

We thank members of the Lingner lab for discussions and reagents; Romain Hamelin, Florence Armand and Diego Chiappe at the Proteomics Core Facility at EPFL for mass spectrometry analysis; the Proteomic Expression Core Facility at EPFL for HEK293E cells; the Flow Cytometry Core Facility at EPFL for technical advice; and the Bioimaging and Optics Platform at EPFL for training and maintenance of the Zeiss-Axioplan microscope. C.-Y.G.L. was supported by an EMBO long-term postdoctoral fellowship and A.C.N. by a PhD fellowship from the Boehringer-Ingelheim Fonds. Research in J.L.’s laboratory was supported by the Swiss National Science Foundation (SNSF), the SNSF-funded National Centre of Competence in Research RNA and Disease Network, an Innovative Training Network (ITN) (aDDRess) from the European Commission’s Seventh Framework Programme (grant No 812829) and EPFL.

## Author contributions

G.L. and J.L. conceived the study initially. C.-Y.G.L., A.C.N., T.L. and A.V. planned and carried out the experiments that are shown in the paper, G.L. constructed the FLAG-tagged cell line and carried out pilot experiments, and C.-Y.G.L. and J.L. wrote the manuscript with contributions from A.C.N.

## Methods

### RESOURCE AVAILABILITY

#### Lead Contact

Further information and requests for resources and reagents should be directed to and will be fulfilled by the Lead Contact, Joachim Lingner.

#### Materials Availability

- Plasmids generated in this study are available upon request.
- This study did not generate new unique reagents.

#### Data and Code Availability

The mass spectrometry proteomics data have been deposited to the ProteomeXchange Consortium via the PRIDE ^65^ partner repository with the dataset identifier PXD018712

Reviewer account details:

**Password:** 0QUxmNza

### EXPERIMENTAL MODEL AND SUBJECT DETAILS

#### Cell lines

- HEK293E cells: A subclone of the <u>H</u>uman <u>E</u>mbryonic <u>K</u>idney (HEK) epithelium suspension cell line expressing the EBNA-1 protein (female origin) was cultured in EX-CELL^(r)^ 293 Serum-Free Medium (Merck, 14571C) containing 4 mM GlutaMAX supplement (TermoFisher, 35050061) at 37°C and 5% CO_2_. Stable cell lines expressing endogenous FLAG-TRF1 and FLAG-TRF2 (heterozygous) were generated by retroviral infection and selection in puromycin-containing medium.
- HeLa-Long cells: A super-telomerase population of HeLa cells (with approximately 33 kb long telomeres) expressing hTERT from the LTR (long terminal repeat) promoter and human telomerase RNA from the U1 promoter generated by retroviral infection^66^. The parental HeLa cell line (female origin) was obtained from ATCC. Cells were cultured in Dulbecco’s Modified Eagle Medium (DMEM) supplemented with 10% Tet System Approved FBS (Clontech), 100 U/ml penicillin and 100 µg/ml streptomycin (Gibco) at 37°C and 5% CO_2_.

## METHOD DETAILS

### CRISPR/Cas9 gene editing and transfections

HEK 293E cl75 was generated by transfecting pSpCas9(BB)-2A-Puro (PX459) plasmid expressing gRNAs targeting the genomic region proximal to the start codon of TRF1 (TRF1-gRNA: 5’-GCGAGCCATTTAACATGGCGG-3’) or TRF2 (TRF2-gRNA: 5’-TCTATCATGGCCGCGGGAGC-3’) and non-linearized repair template plasmid^67^. The repair template contained 600bp homology region flanking both sites of the start codon of TRF1 or TRF2 amplified from genomic DNA of HEK293E cells, and combined with three tandemly repeated FLAG tags, EcoRI restriction site, Kozak sequence and linker base pairs by using overlapping PCR. The PCR fragments were cloned into TOPO Zero Blunt for Sequencing (Invitrogen). Cells were transfected in a 6-well plate containing 2 mL DMEM/FBS, 500 µL OptiMEM and 10 µL Lipofectamine with 3 µg DNA of gRNA sequence and 3 µg repair template. Cells were split the next day and selected with 1µg/mL Puromycin (conc. 1µg/mL, #ant-pr-1, Invivogen) for 4 days. Single-cell clones were obtained by limiting dilution and were screened for the presence of the Flag-Tag by PCR. For screening, genomic DNA extraction was done by Wizard genomic DNA Purification System (Promega). PCR with primers flanking the region of insertion was performed and positive PCRs were digested with EcoRI to confirm homo/heterozygosity of the edited locus. Positive clones with 3xFLAG sequences were further expanded and the PCR product was subcloned into TOPO Zero Blunt for sequencing and genotyping.

### Cell cycle analysis

HEK293E cells were harvested, washed with PBS and centrifuged at 300 g for 3 min at RT. Pellets were fixed by adding dropwise cold 70% EtOH under vortexing to a final concentration of 1 million cells per ml and stored at 4°C overnight. Cells were washed with PBS once and permeabilized in 0.5% Triton X-100 at a final concentration of 10 million cells per ml for 20 min. Cells were washed with PBS twice and were stained with PBS containing DAPI (2µg/ml) incubating for 10 min at 37°C. Cells were transferred to FACS tubes and cell cycle distribution was determined by flow cytometry (LSRII(SORP)).

### QTIP-iPOND

Cells were pulsed with 10 µM EdU for 10 min and were separated into 2 portions. EdU pulse samples were directly crosslinked by adding formaldehyde (FA) (Sigma, F1635) to 1% FA/ml and fixed for 15 min at room temperature (RT). Cells of Thy chase samples were washed with Thy-containing medium (10 µM Thy) and were resuspended in Thy-containing medium. Cells were then incubated for 4 h and crosslinked as mentioned above. After crosslinking, glycine (pH 2.5) was added to a final concentration of 125 mM for 5 min to quench the crosslinking reaction. Cells were then washed by 1x PBS three times. Cells were lysed with lysis buffer (1% SDS, 50 mM Tris-HCl pH 8.0, 10 mM EDTA pH 8.0, cOmplete-EDTA-free protease inhibitor cocktail [Roche]) for 5 min at RT and washed with cold 1x PBS containing 0.5% BSA and then with cold 1x PBS. Pellets were resuspended in Click reaction cocktail (1x PBS pH 7.4, 10 µM Biotin azide, 10 mM sodium ascorbate, 2 mM CuSO_4_) and incubated for 2 h at RT in the dark and kept in the dark for subsequent steps. Pellets were washed once with cold 1x PBS containing 0.5% BSA and twice with cold 1x PBS. Pellets were then resuspended in LB3 buffer (10 mM Tris-HCl pH 8.0, 200 mM NaCl, 1 mM EDTA, 0.5 mM EGTA, 0.5 mM EGTA, 0.1% Na-Deoxycholate, 0.25% Sodium lauroyl sarkosinate, cOmplete-EDTA-free protease inhibitor cocktail [Roche]) at a final concentration of 10 million cells/ml and sonicated for 7 min on ice using a Branson Tip sonicator (30% power, 20 seconds constant pulse, and 20 seconds pause). Cell lysates were centrifuged at 16,800 rcf for 10 min at 4°C to remove insoluble materials. Next, extracts were dialyzed in a dialysis tube (Thermo Scientific, SnakeSkin Dialysis Tubing, 30K MWCO) against IP buffer (50 mM Tris-Cl pH 8.0, 600 mM NaCl, 10 mM EDTA pH 8.0, 0.75% Triton X-100) with 5x of the sample volume twice (5 h and overnight).

ANTI-FLAG M2 Affinity Agarose Gel (Sigma) (1.6 ml slurry beads/ billion cells) was added to the extract for TRF1/TRF2 immunoprecipitation (IP) overnight at 4°C. The next day, beads were washed 5 times with IP buffer. Elution was performed 5 times with IP buffer containing 100 µg/ml of FLAG peptide (Sigma) incubating 30 min on a wheel for each elution. Note that for the M2 only control, the elution was performed with IP buffer containing FLAG peptide without adding Triton X-100. The eluted samples were further concentrated using Centrifugal Filters (Amicon(r) Ultra-15 10K). For the second IP, Dynabeads MyOne Streptavidin C1 beads (Invitrogen(tm)) (1 ml beads/ 5 billion cells) were added for the IP for 20 h at 4°C. The next day, beads were washed twice with IP buffer (QTIP-iPOND) or LB3 buffer (iPOND only), once with 500 mM NaCl and twice with LB3 buffer at RT. Samples were reverse crosslinked using 2x Laemmli buffer (150 mM Tris-HCl pH 6.8, 4% SDS, 100 mM DTT, 20% glycerol, bromophenol blue) for 30 min at 95°C and were loaded on 10% Mini-PROTEAN TGX Precast Protein Gels (BioRad) for MS analysis.

### MS analysis

In-gel digestion and LC-MS/MS analysis were performed by the proteomics core facility at EPFL which followed a previously published protocol with minor modifications^4^. In brief, SDS-PAGE gel lanes were sliced into 8 fractions. Samples were washed twice in 50% ethanol and 50 mM ammonium bicarbonate (AB, Sigma-Aldrich) for 20 min, and dried by vacuum centrifugation. Reduction was then performed with 10 mM dithioerythritol (Merck-Millipore) for 1 h at 56°C. After washing-drying the samples as above described, an alkylation step was performed with 55 mM Iodoacetamide (Sigma-Aldrich) for 45 min at 37°C in the dark. Samples were washed-dried again and digested overnight at 37°C using modified mass spectrometry grade trypsin (Trypsin Gold, Promega) at a concentration of 12.5 ng/µl in 50 mM AB and 10 mM CaCl_2_. Resulting peptides were extracted in 70% ethanol, 5% formic acid (FA, Merck-Millipore) twice for 20 min with permanent shaking. Samples were further dried by vacuum centrifugation and store at −20 °C.

Peptides were desalted on C18 StageTips^68^ and dried by vacuum centrifugation prior to LC-MS/MS injections. Samples were resuspended in 2% acetonitrile (Biosolve), 0.1% FA and nano-flow separations were performed on a Dinoex Ultimate 3000 RSLC nano UPLC system (Thermo Fischer Scientific) on-line connected with a Lumos Orbitrap Mass Spectrometer (Thermo Fischer Scientific). A capillary precolumn (Acclaim Pepmap C18, 3 µm-100AC, 2 cm x 75µm ID) was used for sample trapping and cleaning. Analytical separations were then performed on a 50cm long capillary column (75 µm ID; in-house packed using ReproSil-Pur C18-AQ 1.9 µm silica beads; Dr. Maisch) at 250 nl/min over a 90-min biphasic gradient. Data Dependent mode was used for MS acquisitions (m/z window: 375 to 1’500) where parent ions were detected in the Orbitrap at a resolution of 240’000 (at m/z 200) while daughter ions were detected on the Linear Ion Trap at low resolution (Rapid mode and Max. injection time of 20ms). Only charge states from 2 to 6 were selected for fragmentation using HCD mode with (collision energy value of 30).

Identifications were performed using MaxQuant (version 1.6.2.10)^69,70^ incorporating the Andromeda search engine^71^ against the UniProt human database containing 73,920 sequences ((Release 2019_01) (www.uniprot.org). Streptavidin sequence was added manually. A concatenated decoy database of common contaminant sequences generated by MazzQuant was used to determine the false discovery rate and exclude false positive hits. Both peptide and protein identifications were filtered at 1% FDR relative to hits against the decoy database built by reversing protein sequences. The minimal peptide length was seven amino acids, at least two peptides were required for protein identification and a minimum of two ratio counts was required to quantify proteins. Carbamidomethylation (C) was set as a fixed modification, whereas oxidation (M), acetylation (Protein N-term) and phosphorylation (STY) were considered as variable modifications. LFQ intensities generated by MaxQuant were used for further analysis.

### Dot blot

Reverse crosslinked DNA was purified by NucleoSpin Gel and PCR Clean-up with buffer NTP (Macherey-Nagel) and eluted in 100 µl pre-warmed H_2_O (65°C). Next, DNA was denatured for 10 min at 95°C and chilled on ice for 10 min. DNA was blotted onto a Hybond N+ nylon membrane (GE Healthcare) and an Amersham Protran 0.2 NC nitrocellulose Western blotting membrane (GE Healthcare) using a BioRad dot blot apparatus. The membranes were UV-crosslinked. The nitrocellulose membrane was directly used for BrdU-immunoblotting. For telomere and Alu signal detection, the Hybond N+ nylon membrane was further denatured with 0.8 M NaOH, 0.5 M NaCl and neutralized with 0.5 M Tris-HCl pH 7.0. The membrane was first blocked in Church buffer (1% BSA, 0.5 mM EDTA pH 8.0, 1M phosphoate buffer, 20% SDS) for 1 h at 65°C and incubated with a ^32^P-labeled telomeric probe overnight at 65°C. Then the membrane was washed with 1x SSC containing 0.5% SDS 3 times for 30 min and exposed to a phosphorimager screen overnight. Radioactive signal was detected with Amersham typhoon and the intensity was quantified in AIDA software version 4.06.034. Next, the telomeric probe was removed by washing with boiling 0.1x SSC containing 1% SDS 3 times for 20 min. Then the membrane was prehybridized with Church buffer at 55°C and hybridized with a ^32^P-labeled Alu probe (5’-TGATCCGCCCGCCTCGGCCTCCCAAAGTG-3’) overnight at 55°C, washed with 1x SSC containing 0.5% SDS 3 times for 30 min and exposed to a phosphorimager screen for 2 days. Radioactive signal was again detected with Amersham typhoon and the intensity was quantified in AIDA software version 4.06.034.

### Immunoblots

For protein detection, samples were resuspended in 2x Laemmli buffer and denatured for 5 min at 95°C. Then samples were loaded on 7.5% Mini-PROTEAN TGX Precast Protein Gels (BioRad) and wet-transferred onto Amersham Protran 0.2 NC nitrocellulose Western blotting membranes (GE Healthcare). For BrdU detection, after dot blotting, the membrane was UV-crosslinked and then followed the general steps of immunoblots. Membranes were blocked in 3% BSA in 1x PBS containing 0.1% Tween 20 (1x PBST) for 1 hour at RT and incubated with primary antibodies overnight at 4°C. The next day the membrane was washed with 1x PBS containing 0.1% Tween 20 three times for 15 min. HRP-conjugated secondary antibodies: anti-mouse IgG HRP-conjugated (W4021, Promega) or anti-rabbit IgG HRP-conjugated (W4011, Promega) in combination with ECL spray (Advansta) were used to reveal the signal on a Fusion FX (Vilber) detector. The following antibodies were used: anti-BrdU (555627, BD Pharmingen™, diluted 1:500 in 1x PBST containing 3%BSA); anti-Vinculin (ab129002, Abcam, diluted 1:10,000 in 1x PBST containing 5% milk); anti-TRF1 (sc-6165-R, Santa Cruz, diluted 1:2,000 in 1x PBST containing 3% BSA).

### Calcium phosphate transfection of siRNA

siGENOME SMARTpools from Horizon Discovery at a final concentration of 5.45 nM were used. HeLa-Long cells were transfected in 10 cm dishes at 30-40% confluency using a calcium phosphate precipitation protocol. 0.5 ml transfection mix was prepared containing 125 mM CaCl_2_, 60 nM siRNA and 1x HBSS pH 7.4 (50 mM HEPES, 280 mM NaCl, 1.19 mM Na_2_HPO_4_•2H_2_O, 10 mM KCl), incubated 10 min at RT and added to the cells covered with 5 ml antibiotic-free DMEM/10% FBS. The cells were harvested 72 h post transfection.

### Telomere fluorescence in situ hybridization (FISH) on metaphase spreads

HeLa-Long cells were treated with 50 ng/ml demecolcine for 2 h. Supernatants were collected and the remaining cells were trypsinized. Trypsinized cells were merged with supernatants and spun down for 5 min at 300 rcf. Pellets were resuspended in 0.056 M KCl and swollen for 7 min at 37°C. Swollen cells were spun down for 3 min at 200 rcf and the supernatant was decanted. The cells were resuspended in fixative (75% methanol, 25% acetic acid) and stored at 4°C overnight. To obtain metaphase spreads, the cell suspensions were dropped on glass slides, incubated 1 min at 70°C in a wet chamber and air-dried overnight. For FISH staining, slides were rehydrated in 1x PBS for 5 min, treated with 4% formaldehyde in 1x PBS for 5 min, washed 3 times with 1x PBS for 5 min, and dehydrated with increasing amounts of ethanol (70%, 95%, 100%) for 5 min at each concentration. Air-dried slides were placed on coverslips with 70 µl hybridization mix (10 mM Tris-HCl pH 7.4, 70% formamide, 0.5% blocking reagent, 89 µM Cy3-OO-(CCCTAA)_3_ PNA probe) and denatured at 80°C for 3 min. Hybridization took place for 3 h in a light-protected humified chamber at RT. The slides were washed twice for 15 min in 70% formamide/10 mM Tris-HCl pH 7.4 and three times for 5 min in 0.1 M Tris-HCl pH 7.4/0.15 M NaCl/0.08% Tween-20 with 0.1 µg/ml DAPI in the second wash. The slides were dehydrated with increasing amounts of ethanol as described above, mounted in Vecta shield embedding medium and stored at −20°C. Images were taken with a Zeiss-Axioplan microscope using a 100x objective.

### Quantitative RT-PCR

Cells were directly lysed in 6-well plates and total RNA was isolated using the NucleoSpin RNA isolation kit (Macherey-Nagel) according to the manufacturer’s protocol with 3 DNase treatments. RT-qPCR was performed on a 7900HT Fast Real-Time System (Applied Biosystems) using the Luna Universal OneStep RT-qPCR kit (NEB) according to the manufacturers protocol in a 384-well reaction plate. No-reverse transcription controls were included for each RNA sample for each primer mix and no-template controls for each primer mix. All experiments were performed in two technical replicates. Relative expression levels were calculated using the 2^−ΔΔ Ct^ method. GAPDH was used as a reference gene for normalization.

### BrdU incorporation and ChIP

HEK293E cells were incubated with BrdU for the indicated times and harvested. Cells were fixed for 15 min at room temperature by 1% methanol-free formaldehyde (Thermo Scientific, 28906) in PBS at a final concentration of 10 million cells/ml. Formaldehyde-treated cells were quenched by 125 mM glycine for 5 min and washed three times with 1x PBS. Cells were lysed with lysis buffer (1% SDS, 50 mM Tris-HCl pH 8.0, 10 mM EDTA pH 8.0, cOmplete-EDTA-free protease inhibitor cocktail [Roche]) for 5 min at RT. Chromatin enriched pellets were resuspend in LB3 buffer (10 mM Tris-HCl pH 8.0, 200 mM NaCl, 1 mM EDTA, 0.5 mM EGTA, 0.5 mM EGTA, 0.1% Na-Deoxycholate, 0.25% Sodium lauroyl sarkosinate, cOmplete-EDTA-free protease inhibitor cocktail [Roche]) at the same concentration as above and sonicated for 10 min at 4°C using a Focused-Ultrasonicator (Covaris, E220, duty 5.0%, PIP: 140 W, cycles: 200, amplitude 0, velocity 0, dwell 0, 0.12 x12 mm glass tubes with AFA fiber). Insoluble material was removed by centrifugation at 20,000 g for 15 min at 4°C and extract were diluted by adding 1.25 volumes IP buffer and precleared with 50% slurry Protein G Sepharose 4 Fast Flow beads (GE Healthcare) (blocked with 100 µg/ml BSA for 30 min at RT) for 1 h at 4°C on a wheel. Pre-cleared lysates corresponding to 2 million cells were immunoprecipitated with 30 µl beads of 50% slurry and 4 µg antibody or 5 µl serum overnight at 4°C on a wheel. The next day, beads were washed with 4 wash buffers sequentially at 4°C: Wash 1: 0.1% SDS, 1% Triton, 2 mM EDTA pH 8.0, 300 mM NaCl; Wash 2: 0.1% SDS, 1% Triton, 2 mM EDTA pH 8.0, 500 mM NaCl; Wash 3: 250 mM LiCl, 1% NP-40, 1 mM EDTA pH 8.0, 10 mM Tris-Cl pH 8.0; Wash 4: 1 mM EDTA pH 8.0 and 10 mM Tris-Cl pH 8.0. Wash 4 was repeated before the beads were incubated in 100 µl crosslink reversal buffer (1% SDS, 0.1 M NaHCO_3_, 0.5 mM EDTA pH 8.0, 20 mM Tris-Cl pH 8.0, 200 µg/ml RNase [DNase-free, Roche]) overnight at 65°C. DNA was stored at −20°C. The following antibodies were used: anti-TRF1 (Rabbit serum 474), anti-POT1 (Rabbit serum 913), RPA32 (Bethyl, A300-244A) and Rabbit IgG (ab172730, Abcam).

### Quantification and statistical analysis

For MS data analysis, the MaxQuant output table “proteinGroups.txt” were processed to assess the significance of outlier ratios using Perseus^72^ and home-made programs written in Perl (v5.18.2) and RStudio (Version 1.2.1335). 1,732 nuclear proteins were first extracted according to annotations from Gene Ontology^57^ and The Human Protein Altas^73^, which contained 22,481 reference proteins that have been shown to localize in nucleus or nucleolus. The ratio between two datasets was normalized to the LFQ intensity value of TRF1 and was then calculated by dividing the number of LFQ intensity values. Only proteins identified in at least 3 of the 4 datasets were included in the statistical analysis in Perseus using a two-tailed t-test with Benjamini-Hochberg FDR correction^74^ which thresholds of 0.05, 0.01, 0.005 and 0.001 on the adjusted p-value and a minimal fold change of 3. Graphical display was made by RStudio. Other statistical analyses in this study were done by unpaired t-test using Prism 8 as specified in figure legends and the results section.

## Supplemental Information

**Table S1 Protein list of QTIP-iPOND**.

**Table S2 Protein list of iPOND**.

**Table S3 Protein list of QTIP**.

**Figure S1** a. Experimental scheme of QTIP-iPOND Thy chase. After 10 min EdU incubation, EdU was washed out and was replaced by medium with thymidine and incubated for 4 hours. The following steps were the same as for QTIP-iPOND EdU pulse. b. Cell cycle profiles of HEK293E Clone 75 cells used for the full-scaled experiment. EdU: 3 biological replicates; Thy: 2 biological replicates. Data are represented as mean ± SD.

**Figure S2 Protein distribution of QTIP-iPOND**. a. EdU pulse over Thy experiment of EdU positive samples. b. EdU pulse over Thy experiment of EdU negative samples. Difference: log_2_(Fold change); Significance: −log(p-value); Blue circles: The 142 proteins with FDR ≤0.001; Orange circles: core replication proteins; Dark green circles: shelterin components; Black curves: FDR=0.05; Violet curves: FDR=0.1; s0 = 3.

**Figure S3 Proteomic analysis of QTIP-iPOND**. a. Comparison of enriched proteins between QTIP-iPOND and iPOND. FDRs <0.05 and s0 = 3 (QTIP-iPOND) and = 0.5 (iPOND). b. The enrichment of histone proteins in iPOND EdU pulse versus Thy chase. EdU pulse and Thy chase were normalized by the fold change of histone H2B.

**Figure S4 Telomere fragility score in the first replicate of the siRNA screen**. The respective score is indicated in brackets. Positive controls: NONO, BLM, WRN and SAMHD1; negative control: SFPQ.

**Figure S5 Confirmation of depletions in the siRNA screen for telomere fragility**. a. Representative image of Western blot for the positive control siTRF1. b. Relative mRNA levels (normalized to siCTL) determined by RT-qPCR for the second and third replicate of the siRNA screen. n=2 (except for siTRF1 with n=11). Data are represented as mean ± SD.

**Figure S6 The BrdU amount of POT1-ChIP in 10-min BrdU incorporation is at the same level as the IgG control**. Result was obtained in triplicate. Data are represented as mean ± SD.

## Reference

1. de Lange, T. Shelterin-Mediated Telomere Protection. Annu. Rev. Genet. 52, 223–247 (2018).

2. Bartocci, C. et al. Isolation of Chromatin from Dysfunctional Telomeres Reveals an Important Role for Ring1b in NHEJ-Mediated Chromosome Fusions. Cell Rep. 7, 1320–1332 (2014).

3. Déjardin, J. & Kingston, R. E. Purification of Proteins Associated with Specific Genomic Loci. Cell 136, 175–186 (2009).

4. Grolimund, L. et al. A quantitative telomeric chromatin isolation protocol identifies different telomeric states. Nat. Commun. 4, (2013).

5. Fagagna, F. d’Adda di et al. A DNA damage checkpoint response in telomere-initiated senescence. Nature 426, 194–198 (2003).

6. Maciejowski, J. & de Lange, T. Telomeres in cancer: tumour suppression and genome instability. Nat. Rev. Mol. Cell Biol. 18, 175–186 (2017).

7. Armanios, M. Syndromes of Telomere Shortening. Annu. Rev. Genomics Hum. Genet. 10, 45–61 (2009).

8. Glousker, G. & Lingner, J. When Telomerase Causes Telomere Loss. Dev. Cell 44, 281–283 (2018).

9. Glousker, G., Touzot, F., Revy, P., Tzfati, Y. & Savage, S. A. Unraveling the pathogenesis of Hoyeraal-Hreidarsson syndrome, a complex telomere biology disorder. Br. J. Haematol. 170, 457–471 (2015).

10. Kong, C. M., Lee, X. W. & Wang, X. Telomere shortening in human diseases. FEBS J. 280, 3180–3193 (2013).

11. Sarek, G., Marzec, P., Margalef, P. & Boulton, S. J. Molecular basis of telomere dysfunction in human genetic diseases. Nat. Struct. Mol. Biol. 22, 867–874 (2015).

12. Crabbe, L. Defective Telomere Lagging Strand Synthesis in Cells Lacking WRN Helicase Activity. Science 306, 1951–1953 (2004).

13. Deng, Z. et al. Inherited mutations in the helicase RTEL1 cause telomere dysfunction and Hoyeraal–Hreidarsson syndrome. Proc. Natl. Acad. Sci. 110, E3408–E3416 (2013).

14. Drosopoulos, W. C., Kosiyatrakul, S. T. & Schildkraut, C. L. BLM helicase facilitates telomere replication during leading strand synthesis of telomeres. J. Cell Biol. 210, 191–208 (2015).

15. Rhodes, D. & Lipps, H. J. G-quadruplexes and their regulatory roles in biology. Nucleic Acids Res. 43, 8627–8637 (2015).

16. Griffith, J. D. et al. Mammalian telomeres end in a large duplex loop. Cell 97, 503–514 (1999).

17. Sarek, G., Vannier, J.-B., Panier, S., Petrini, J. H. J. & Boulton, S. J. TRF2 Recruits RTEL1 to Telomeres in S Phase to Promote T-Loop Unwinding. Mol. Cell 57, 622–635 (2015).

18. Uringa, E.-J. et al. RTEL1 contributes to DNA replication and repair and telomere maintenance. Mol. Biol. Cell 23, 2782–2792 (2012).

19. Vannier, J.-B., Pavicic-Kaltenbrunner, V., Petalcorin, M. I. R., Ding, H. & Boulton, S. J. RTEL1 Dismantles T Loops and Counteracts Telomeric G4-DNA to Maintain Telomere Integrity. Cell 149, 795–806 (2012).

20. Azzalin, C. M. & Lingner, J. Telomere functions grounding on TERRA firma. Trends Cell Biol. 25, 29–36 (2015).

21. Azzalin, C. M., Reichenbach, P., Khoriauli, L., Giulotto, E. & Lingner, J. Telomeric Repeat Containing RNA and RNA Surveillance Factors at Mammalian Chromosome Ends. Science 318, 798–801 (2007).

22. Feretzaki, M., Renck Nunes, P. & Lingner, J. Expression and differential regulation of human TERRA at several chromosome ends. RNA 25, 1470–1480 (2019).

23. Porro, A. et al. Functional characterization of the TERRA transcriptome at damaged telomeres. Nat. Commun. 5, 5379 (2014).

24. Arora, R. et al. RNaseH1 regulates TERRA-telomeric DNA hybrids and telomere maintenance in ALT tumour cells. Nat. Commun. 5, 5220 (2014).

25. Balk, B. et al. Telomeric RNA-DNA hybrids affect telomere-length dynamics and senescence. Nat. Struct. Mol. Biol. 20, 1199–1205 (2013).

26. Luke, B. et al. The Rat1p 511 to 311 Exonuclease Degrades Telomeric Repeat-Containing RNA and Promotes Telomere Elongation in Saccharomyces cerevisiae. Mol. Cell 32, 465–477 (2008).

27. Petti, E. et al. SFPQ and NONO suppress RNA:DNA-hybrid-related telomere instability. Nat. Commun. 10, 1001 (2019).

28. Pfeiffer, V., Crittin, J., Grolimund, L. & Lingner, J. The THO complex component Thp2 counteracts telomeric R-loops and telomere shortening. EMBO J. 32, 2861–2871 (2013).

29. Sfeir, A. et al. Mammalian Telomeres Resemble Fragile Sites and Require TRF1 for Efficient Replication. Cell 138, 90–103 (2009).

30. Majerská, J., Redon, S. & Lingner, J. Quantitative telomeric chromatin isolation protocol for human cells. Methods 114, 28–38 (2017).

31. Sirbu, B. M., Couch, F. B. & Cortez, D. Monitoring the spatiotemporal dynamics of proteins at replication forks and in assembled chromatin using isolation of proteins on nascent DNA. Nat. Protoc. 7, 594–605 (2012).

32. Sirbu, B. M. et al. Identification of Proteins at Active, Stalled, and Collapsed Replication Forks Using Isolation of Proteins on Nascent DNA (iPOND) Coupled with Mass Spectrometry. J. Biol. Chem. 288, 31458–31467 (2013).

33. Cortez, D. Proteomic Analyses of the Eukaryotic Replication Machinery. in Methods in Enzymology vol. 591 33–53 (Elsevier, 2017).

34. Somyajit, K. et al. Redox-sensitive alteration of replisome architecture safeguards genome integrity. Science 358, 797–802 (2017).

35. Wessel, S. R., Mohni, K. N., Luzwick, J. W., Dungrawala, H. & Cortez, D. Functional Analysis of the Replication Fork Proteome Identifies BET Proteins as PCNA Regulators. Cell Rep. 28, 3497-3509.e4 (2019).

36. Li, F. et al. The BUB3-BUB1 Complex Promotes Telomere DNA Replication. Mol. Cell 70, 395-407.e4 (2018).

37. Majerska, J., Feretzaki, M., Glousker, G. & Lingner, J. Transformation-induced stress at telomeres is counteracted through changes in the telomeric proteome including SAMHD1. Life Sci. Alliance 1, e201800121 (2018).

38. Lopez-Contreras, A. J. et al. A Proteomic Characterization of Factors Enriched at Nascent DNA Molecules. Cell Rep. 3, 1105–1116 (2013).

39. Galati, A. et al. TRF2 Controls Telomeric Nucleosome Organization in a Cell Cycle Phase-Dependent Manner. PLoS ONE 7, e34386 (2012).

40. Tommerup, H., Dousmanis, A. & de Lange, T. Unusual chromatin in human telomeres. Mol. Cell. Biol. 14, 5777–5785 (1994).

41. Murga, M. et al. Global chromatin compaction limits the strength of the DNA damage response. J. Cell Biol. 178, 1101–1108 (2007).

42. Greider, C. W. Regulating telomere length from the inside out: the replication fork model. Genes Dev. 30, 1483–1491 (2016).

43. Miller, K. M., Rog, O. & Cooper, J. P. Semi-conservative DNA replication through telomeres requires Taz1. Nature 440, 824–828 (2006).

44. Zeng, X. et al. Administration of a Nucleoside Analog Promotes Cancer Cell Death in a Telomerase-Dependent Manner. Cell Rep. 23, 3031–3041 (2018).

45. Benyelles, M. et al. Impaired telomere integrity and rRNA biogenesis in PARN-deficient patients and knock-out models. EMBO Mol. Med. 11, (2019).

46. Moon, D. H. et al. Poly(A)-specific ribonuclease (PARN) mediates 3’-end maturation of the telomerase RNA component. Nat. Genet. 47, 1482–1488 (2015).

47. Tummala, H. et al. Poly(A)-specific ribonuclease deficiency impacts telomere biology and causes dyskeratosis congenita. J. Clin. Invest. 125, 2151–2160 (2015).

48. Yuan, F., Li, G. & Tong, T. Nucleolar and coiled-body phosphoprotein 1 (NOLC1) regulates the nucleolar retention of TRF2. Cell Death Discov. 3, 17043 (2017).

49. Meng, L., Hsu, J. K., Zhu, Q., Lin, T. & Tsai, R. Y. L. Nucleostemin inhibits TRF1 dimerization and shortens its dynamic association with the telomere. J. Cell Sci. 124, 3706–3714 (2011).

50. Zhu, Q., Yasumoto, H. & Tsai, R. Y. L. Nucleostemin delays cellular senescence and negatively regulates TRF1 protein stability. Mol. Cell. Biol. 26, 9279–9290 (2006).

51. Zhu, Q. et al. GNL3L stabilizes the TRF1 complex and promotes mitotic transition. J. Cell Biol. 185, 827–839 (2009).

52. Yoo, J. E., Oh, B.-K. & Park, Y. N. Human PinX1 Mediates TRF1 Accumulation in Nucleolus and Enhances TRF1 Binding to Telomeres. J. Mol. Biol. 388, 928–940 (2009).

53. Schmidt, J. C., Zaug, A. J. & Cech, T. R. Live Cell Imaging Reveals the Dynamics of Telomerase Recruitment to Telomeres. Cell 166, 1188-1197.e9 (2016).

54. Jády, B. E., Richard, P., Bertrand, E. & Kiss, T. Cell Cycle-dependent Recruitment of Telomerase RNA and Cajal Bodies to Human Telomeres. Mol. Biol. Cell 17, 944–954 (2006).

55. Tomlinson, R. L., Ziegler, T. D., Supakorndej, T., Terns, R. M. & Terns, M. P. Cell cycle-regulated trafficking of human telomerase to telomeres. Mol. Biol. Cell 17, 955–965 (2006).

56. Zhou, X. Z. & Lu, K. P. The Pin2/TRF1-Interacting Protein PinX1 Is a Potent Telomerase Inhibitor. Cell 107, 347–359 (2001).

57. Ashburner, M. et al. Gene Ontology: tool for the unification of biology. Nat. Genet. 25, 25–29 (2000).

58. The Gene Ontology Consortium. The Gene Ontology Resource: 20 years and still GOing strong. Nucleic Acids Res. 47, D330–D338 (2019).

59. Brockers, K. & Schneider, R. Histone H1, the forgotten histone. Epigenomics 11, 363–366 (2019).

60. Andreyeva, E. N. et al. Regulatory functions and chromatin loading dynamics of linker histone H1 during endoreplication in Drosophila. Genes Dev. 31, 603–616 (2017).

61. Thiriet, C. & Hayes, J. J. Linker Histone Phosphorylation Regulates Global Timing of Replication Origin Firing. J. Biol. Chem. 284, 2823–2829 (2009).

62. Pinzaru, A. M. et al. Telomere Replication Stress Induced by POT1 Inactivation Accelerates Tumorigenesis. Cell Rep. 15, 2170–2184 (2016).

63. Flynn, R. L. et al. TERRA and hnRNPA1 orchestrate an RPA-to-POT1 switch on telomeric single-stranded DNA. Nature 471, 532–536 (2011).

64. Glousker, G., Briod, A.-S., Quadroni, M. & Lingner, J. Human POT1 Prevents Severe Telomere Instability Induced by Homology Directed DNA Repair. http://biorxiv.org/lookup/doi/10.1101/2020.01.20.912642 (2020) doi:10.1101/2020.01.20.912642.

65. Perez-Riverol, Y. et al. The PRIDE database and related tools and resources in 2019: improving support for quantification data. Nucleic Acids Res. 47, D442–D450 (2019).

66. Cristofari, G. & Lingner, J. Telomere length homeostasis requires that telomerase levels are limiting. EMBO J. 25, 565–574 (2006).

67. Briod, A.-S., Glousker, G. & Lingner, J. RADX Sustains POT1 Function at Telomeres to Counteract RAD51 Binding, which Triggers Telomere Fragility. http://biorxiv.org/lookup/doi/10.1101/2020.01.20.912634 (2020) doi:10.1101/2020.01.20.912634.

68. Rappsilber, J., Mann, M. & Ishihama, Y. Protocol for micro-purification, enrichment, pre-fractionation and storage of peptides for proteomics using StageTips. Nat. Protoc. 2, 1896–1906 (2007).

69. Cox, J. & Mann, M. MaxQuant enables high peptide identification rates, individualized p.p.b.-range mass accuracies and proteome-wide protein quantification. Nat. Biotechnol. 26, 1367–1372 (2008).

70. Cox, J., Michalski, A. & Mann, M. Software Lock Mass by Two-Dimensional Minimization of Peptide Mass Errors. J. Am. Soc. Mass Spectrom. 22, 1373–1380 (2011).

71. Cox, J. et al. Andromeda: A Peptide Search Engine Integrated into the MaxQuant Environment. J. Proteome Res. 10, 1794–1805 (2011).

72. Tyanova, S. et al. The Perseus computational platform for comprehensive analysis of (prote)omics data. Nat. Methods 13, 731–740 (2016).

73. Uhlen, M. et al. Tissue-based map of the human proteome. Science 347, 1260419–1260419 (2015).

74. Benjamini, Y. & Hochberg, Y. Controlling the False Discovery Rate: A Practical and Powerful Approach to Multiple Testing. J. R. Stat. Soc. Ser. B Methodol. 57, 289–300 (1995).

