## Supplementary material for "The Human Telomeric Proteome During Telomere Replication": figS3.pdf

Figure S3

**a**

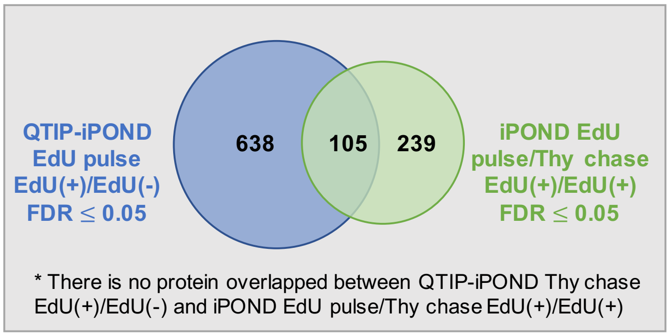


**b**

| Protein name | Gene name | iPOND EdU/Thy (EdU+/EdU+) | |
|---|---|---|---|
| | | Avg. FC | $Log_2$ (FC) of 4 replicates |
| Histone H1.0 | H1F0 | 0.5 | |
| Histone H1.2; Histone H1.1 | HIST1H1C/A | 0.7 | |
| Histone H1.3 | HIST1H1D | 0.6 | |
| Histone H1.4 | HIST1H1E | 0.6 | |
| Histone H1.5 | HIST1H1B | 0.4 | |
| Histone H1x | H1FX | 0.9 | |
| Histone H2A | HIST1H2AB | 1.4 | |
| Core histone macro-H2A.1; Histone H2A | H2AFY | 0.8 | |
| Core histone macro-H2A.2; Histone H2A | H2AFY2 | 0.8 | |
| Histone H2A.V; Histone H2A.Z; Histone H2A | H2AFV;H2AFZ | 0.2 | |
| Histone H2A.X | H2AFX | 1.0 | |
| Histone H2B | HIST1H2BN | 1.0 | |
| Histone H3 | HIST2H3PS2 | 0.8 | |
| Histone H3.1; Histone H3.1t | HIST1H3A;HIST3H3 | 0.8 | |
| Histone H3; Histone H3.3; Histone H3.3C | H3F3B;H3F3A;H3F3C | 2.0 | |
| Histone H4 | HIST1H4A | 1.9 | |

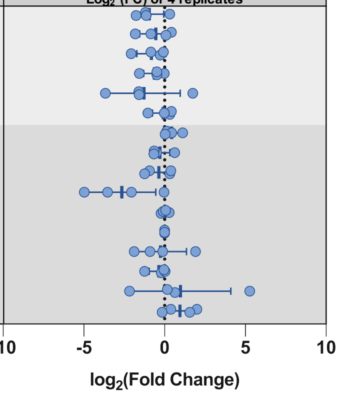
