## Supplementary material for "The Human Telomeric Proteome During Telomere Replication": FigS4.pdf

Figure S4

| Score ≥ 0.4 | 0.2 ≤ Score < 0.4 | Score < 0.2 |
|---|---|---|
| NONO (1.22) | TCEA1 (0.37) | CCDC86 (0.19) |
| NLE1 (1.07) | GPKOW (0.36) | SFPQ (0.11) |
| HMGN5 (1.03) | DDX55 (0.35) | NOC3L (0.08) |
| SETD7 (1.03) | C7orf50 (0.29) | RTF1 (0.04) |
| BLM (0.85) | RNPS1 (0.26) | DAXX (0.04) |
| PAF1 (0.81) | | NCL (0.00) |
| CXorf56 (0.71) | | MAD1L1 (0.00) |
| CENPH (0.66) | | |
| RCC1 (0.63) | | |
| HMGB2 (0.61) | | |
| SPTY2D1 (0.60) | | |
| REXO4 (0.57) | | |
| PHF2 (0.56) | | |
| SAMHD1 (0.49) | | |
| C1orf131 (0.45) | | |
| WRN (0.41) | | |
| LARP7 (0.40) | | |
