## Supplementary figures and images for "The Human Telomeric Proteome During Telomere Replication"

### FigS1.pdf

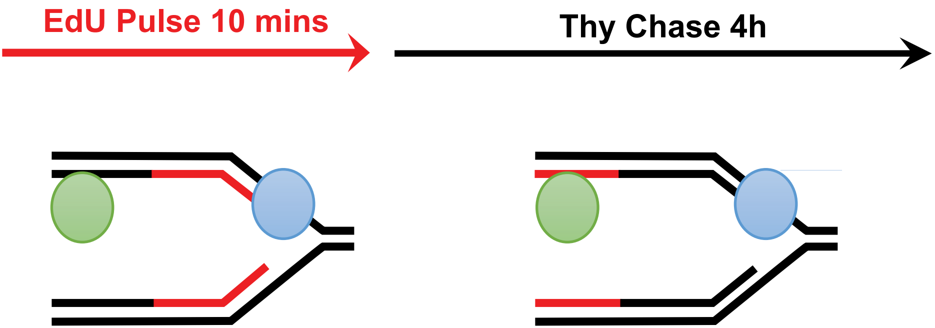

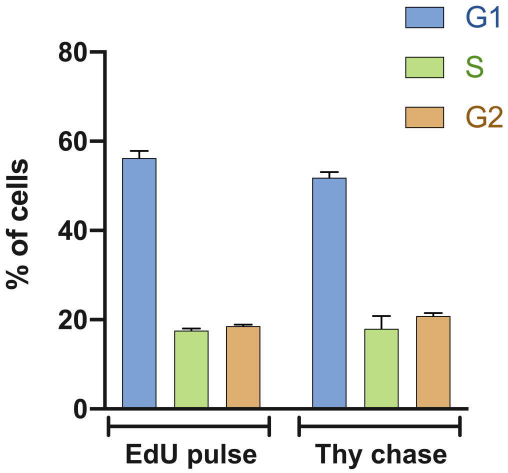

### FigS2.pdf

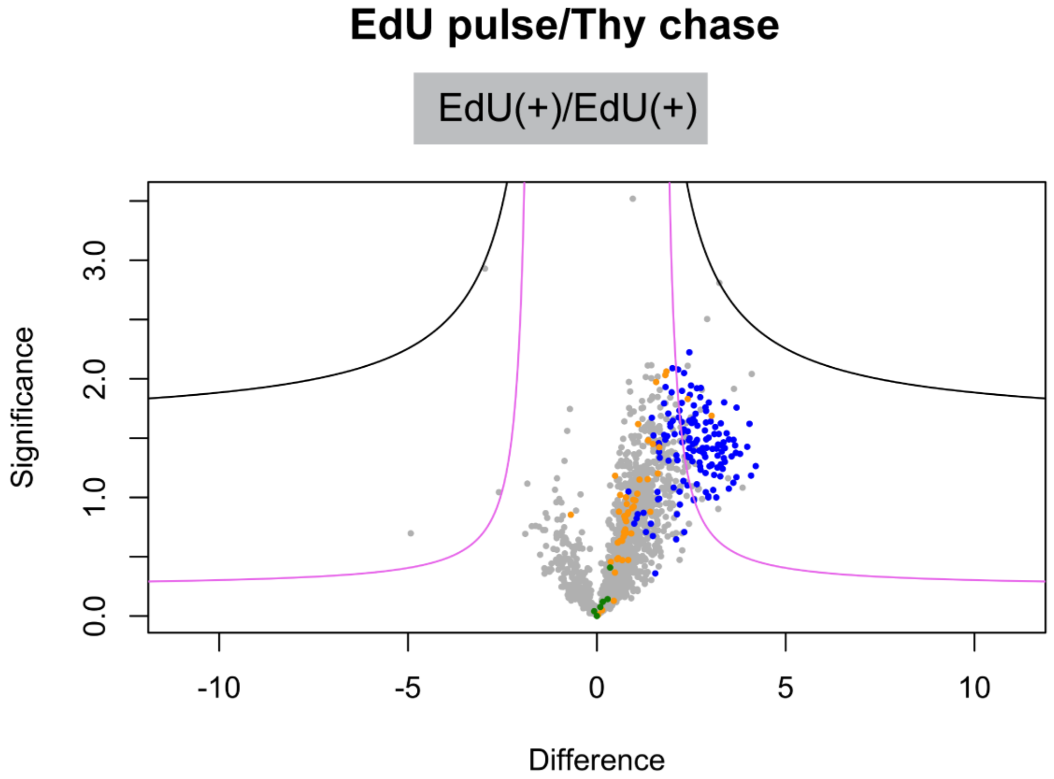

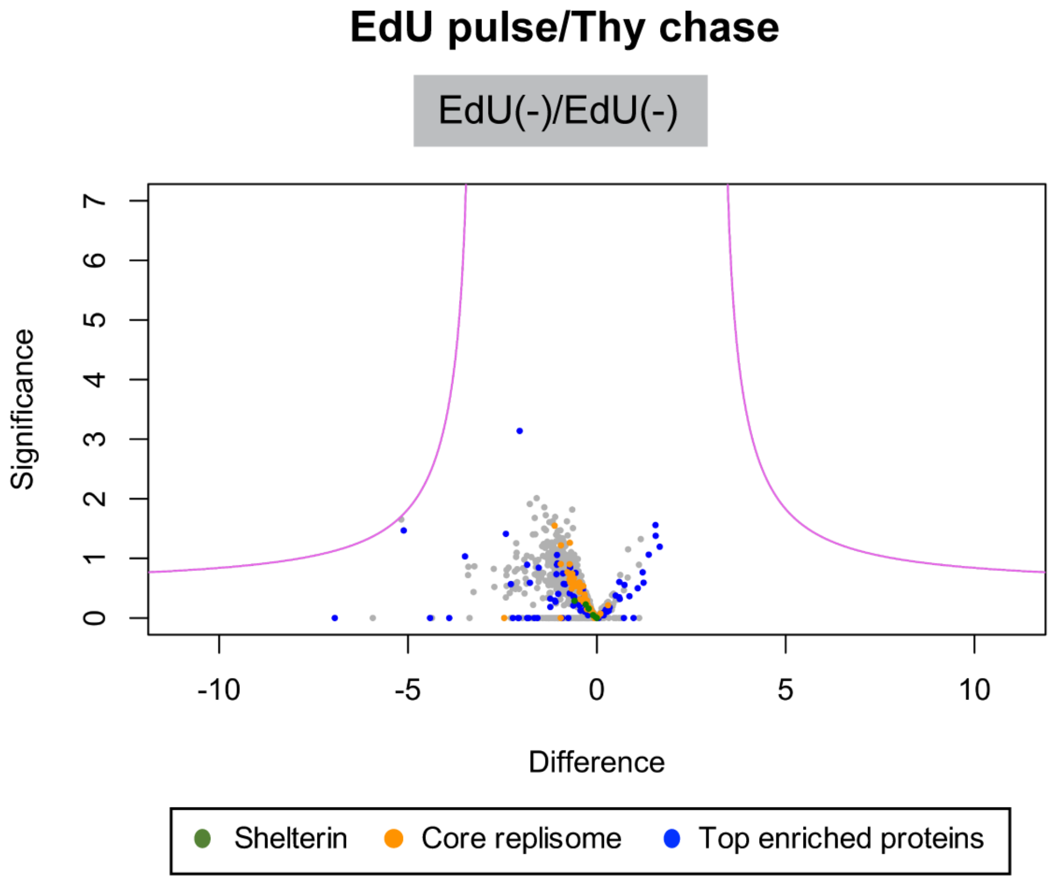

### FigS5.pdf

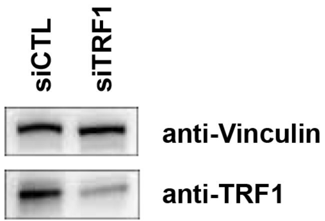

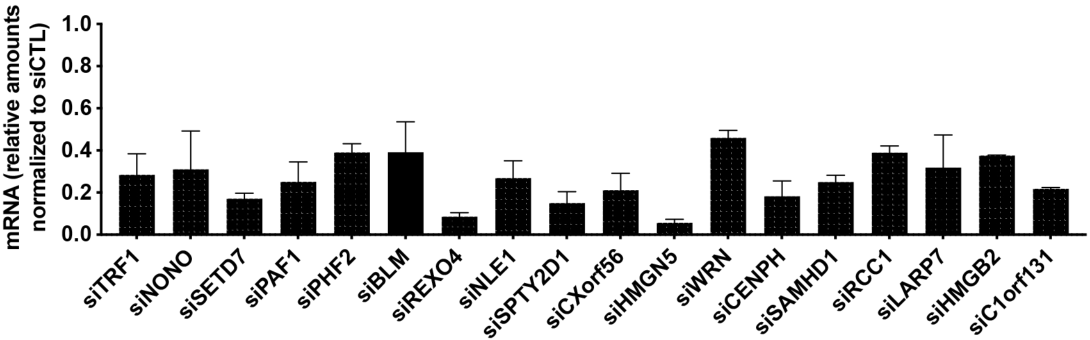

### FigS6.pdf

Figure S6

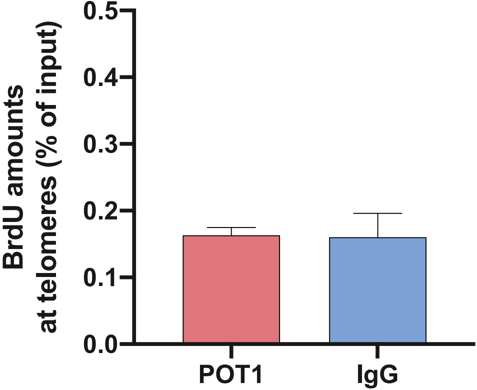
